# Quantitative proteomic analysis reveals apoE4-dependent phosphorylation of the actin-regulating protein VASP

**DOI:** 10.1101/2022.06.06.495052

**Authors:** Zeynep Cakir, Samuel J. Lord, Yuan Zhou, Gwendolyn M. Jang, Benjamin J. Polacco, Manon Eckhardt, David Jimenez-Morales, Billy W. Newton, Adam L. Orr, Jeffrey R. Johnson, Alexandre da Cruz, R. Dyche Mullins, Nevan J. Krogan, Robert W. Mahley, Danielle L. Swaney

## Abstract

Apolipoprotein (apo) E4 is the major genetic risk factor for Alzheimer’s Disease (AD). While neurons generally produce a minority of the apoE in the central nervous system, neuronal expression of apoE increases dramatically in response to stress and is sufficient to drive pathology. Currently, the molecular mechanisms of how apoE4 expression may regulate pathology are not fully understood. Here we expand upon our previous studies measuring the impact of apoE4 on protein abundance to include the analysis of protein phosphorylation and ubiquitylation signaling in isogenic Neuro-2a cells expressing apoE3 or apoE4. ApoE4 expression resulted in a dramatic increase in VASP S235 phosphorylation in a PKA-dependent manner. This phosphorylation disrupted VASP interactions with numerous actin cytoskeletal and microtubular proteins. Reduction of VASP S235 phosphorylation via PKA inhibition resulted in a significant increase in filopodia formation and neurite outgrowth in apoE4-expressing cells, exceeding levels observed in apoE3-expressing cells. Our results highlight the pronounced and diverse impact of apoE4 on multiple modes of protein regulation and identify protein targets to restore apoE4-related cytoskeletal defects.

## Introduction

Apolipoprotein (apo) E is the major cholesterol transport protein in the brain and exists as three allelic variants, apoE2, apoE3, and apoE4, which each differ by single amino acid interchanges (1–3). The apoE4 variant is the major genetic risk factor for Alzheimer’s disease (AD) (4–6), occuring in 60–70% of all sporadic and familial AD cases (2, 4–6). ApoE4 lowers the age of onset and increases the risk of developing AD, and considerable evidence indicates that apoE4 has a fundamental role in AD neurodegeneration (2, 3, 7–15). However, the precise molecular mechanisms by which apoE4 drives pathology remain unclear (16, 17).

Current evidence suggests that neuronal expression of apoE4 is important to this process. Though neurons produce a minority of the apoE in the central nervous system (~20%) (18), neuronal apoE expression levels increase dramatically in response to stress via a neuron-specific splicing mechanism (19). Furthermore, neuronal expression of apoE4 is sufficient to drive pathology (2, 10, 13, 18, 20), likely via an apoE4 gain-of-function mechanism as compared to apoE3 (21).

Numerous studies have shown that apoE4-driven neuropathology is associated with a variety of cellular phenotypes, including mitochondrial dysfunction (22–24), cytoskeletal defects and altered microtubule structure (13). Remarkably, apoE4 expression increases phosphorylation of the microtubule-associated protein tau by activating the Erk pathway in the hippocampus of Neuron-Specific Enolase (NSE)-apoE transgenic mice and in cultured neuronal cells (25). Furthermore, apoE4 induces neurofibrillary tangle (NFT) structures that include phosphorylated tau protein (26). Treatment of cultured neuronal cells with apoE4-lipid complexes impairs neurite outgrowth compared to apoE3-lipid complexes (27–29). Likewise, neuroblastoma cells stably expressing apoE4 display impaired neurite outgrowth; however, this can be restored by site-directed mutagenesis of apoE4 or by treatment with small molecule structure correctors that convert the apoE4 structure to one that resembles apoE3 (30, 31).

Unbiased global proteomic analysis of protein abundance and post-translational modifications (PTMs) by mass spectrometry (MS) represents a powerful mechanism to discover regulatory biological mechanisms. However, to date, efforts to identify the proteomic impact of apoE4 by MS have primarily been focused on specific organelles or subcompartments (32, 33), and have identified few differentially regulated proteins. Recently, we reported a global protein abundance analysis of Neuro-2a cells expressing apoE3 or apoE4 revealing hundreds of differentially regulated proteins and correlating these differences to apoE4-specific cellular processes (24). To further our understanding of the molecular mechanisms that may regulate apoE4-specific pathology, we have expanded upon our previous studies to include the analysis of protein phosphorylation and ubiquitylation signaling in isogenic Neuro-2a cells expressing human apoE3 or apoE4. The results highlight the diverse impact of apoE4 on multiple modes of protein regulation and identify a mechanism of apoE4-mediated cytoskeletal biology.

### Experimental Procedures

#### Experimental Design and Statistical Rationale

Global proteomics analyses measuring protein abundances, phosphorylated peptides, and ubiquitylated peptides were each performed in biological triplicate for both the apoE3 and apoE4 Neuro-2a cells. Each sample was analyzed on the mass spectrometer in technical duplicate. APMS experiments were performed in biological triplicate for each condition, with one replicate of data collected on the mass spectrometer. Label-free quantification and statistical analysis for both global proteomics and APMS data was performed using MSstats and resulting Log2 fold changes and adjusted p-values are reported (34).

#### Cell culture

Mouse neuroblastoma Neuro-2a cell lines stably expressing full-length human apoE3 or apoE4 have been described previously (25). Cells were cultured in MEM medium containing 1x glutamax (Cat.#41090036, Gibco), 10% fetal bovine serum (Cat.#A3160402, Gibco), 1% nonessential amino acids (Cat.#11140050, Gibco), 1 mM sodium pyruvate (Cat.#11360-070, Gibco).

#### Cell lysis and digestion

Nearly confluent Neuro-2a apoE3 or apoE4 cells growing on 15cm dishes were rinsed once with 10 mL of ice-cold 1x PBS and scraped off the plate in 10 mL fresh ice-cold 1x PBS. Cells were pelleted at 4°C for 10 min at 1,500 rpm, the supernatant was decanted, and cells rinsed by resuspending in 10 mL fresh ice cold 1x PBS and pelleting again at 4°C for 10 min at 1,500 rpm. Finally, cells were flash frozen on dry ice and stored at –80°C until all replicates were collected. Three independent sets of cultures were harvested separately.

Cells were lysed by probe sonication on ice in a buffer composed of 8 M urea, 100 mM ammonium bicarbonate pH 8.0, 150 mM NaCl, and cOmplete mini protease (Cat. #1183615300, Roche) and phosphatase inhibitors (PHOSS-RO, Roche). Proteins were reduced by the addition of 4 mM TCEP and incubated at 22°C for 30 min. Cysteines were alkylated by the addition of 10 mM iodoacetamide and lysates were incubated in the dark at 22°C for 30 min. Alkylation was quenched by the addition of 10 mM DTT and lysates were further incubated in the dark at 22°C for 30 min. Next, proteins were digested overnight at 37°C by the addition of trypsin (Promega) at a 1:100 enzyme:substrate ratio. Samples were acidified to pH 2 with trifluoroacetic acid and desalted on C18 SepPak cartridges (Waters).

#### Phosphopeptide enrichment

Immobilized metal affinity chromatography (IMAC) beads for phosphopeptide enrichment were prepared by stripping Ni-NTA Superflow beads (Qiagen) with 100 mM EDTA (pH 7.5), followed by loading of the beads with iron by incubated in 15 mM ferric chloride solution. The beads were then washed with water and then 15uL of beads were loaded into the top of a C18 microspin column. Approximately 1mg of desalted peptides was resuspended in 75% acetonitrile, 0.15% trifluoroacetic acid, and loaded onto IMAC tips by pipetting and incubated for 4 min, with intermittent mixing by pipetting. The lysate was then passed through the column, followed by 4 washes with 200 μL of 80% acetonitrile, 0.1% trifluoroacetic acid. The column was then equilibrated twice for 2 min with 200 μL of 0.5% formic acid, with intermittent mixing by pipetting. Next, the bound phosphorylated peptides were eluted from the IMAC beads onto the C18 column by the addition of 200 μL of 500 mM potassium phosphate buffer (pH 7), which was incubated with the beads for 3 min with intermittent mixing by pipetting. The potassium phosphate elution was repeated once, and then the column was equilibrated twice for 15 s with 200 μL of 0.5% formic acid. Finally, the phosphopeptides were eluted from the C18 column twice by the addition of 75 μL of 50% acetonitrile in 0.1% formic acid and the eluted phosphopeptides were dried by vacuum centrifugation.

#### Ubiquitylated peptide enrichment

Approximately 1/8th of a tube of anti PTMScan Ubiquitin Remnant Motif antibody beads (Cell Signaling Technologies) was used per sample. All steps were performed at 4°C unless otherwise noted. The beads were first washed twice in IAP buffer (50 mM MOPS, 10 mM sodium phosphate buffer (pH 7.2), 50 mM NaCl) and then ~5 mg of desalted peptides that had been resuspended in IAP was added to the beads. The samples were incubated with rotation for 90 min, and then the beads were washed twice with IAP buffer, followed by two washes with water. Next, peptides were eluted from the beads by the addition of 60 μL of 0.15% trifluoroacetic acid to the beads for 10 min with gentle vortexing. The supernatant containing the ubiquitin enriched peptide eluent was then transferred to a new tube, and the elution was repeated by the addition of another 60 μL of 0.15% trifluoroacetic acid. The second elution was combined with the first, and then desalted on C18 columns (The Nest Group), and the desalted peptides were dried by vacuum centrifugation.

#### Mass spectrometry data acquisition

Peptides were resuspended in 4% formic acid and 3% acetonitrile and directly injected on a 75 μm ID column (New Objective) packed with 25 cm of Reprosil C18 1.9 μm, 120 Å particles (Dr. Maisch) for AB, PH, and UB experiments. For APMS experiments, peptides were resuspended in 0.1% formic and directly injected onto a PepSep column (Bruker) packed with 15 cm of Reprosil C18 1.5 μm, 120 Å particles (Dr. Maisch). Peptides were eluted in positive ion mode into an Orbitrap Fusion Tribrid mass spectrometer (Thermo Fisher Scientific) with an acetonitrile gradient delivered by an Easy1000 nLC system (PH, AB, and UB) or an Orbitrap Fusion Lumos Tribrid mass spectrometer (Thermo Fisher Scientific) with an acetonitrile gradient delivered by an Easy1200 nLC system (APMS) (Thermo Fisher Scientific). All mobile phases contained 0.1% formic acid as buffer A. For AB and PH experiments the total MS acquisition time was 180 min, during which mobile phase buffer B (0.1% formic acid in 90% acetonitrile) was ramped from 5% to 16% B over 114 min, followed by a ramp to 30% buffer B over a subsequent 50 min, and then a ramp to 95% B to wash the column. For the UB experiment the total MS acquisition time was 120 min, during which mobile phase buffer B (0.1% formic acid in 90% acetonitrile) was ramped from 5% to 33% B over 105 min, followed by a ramp to 100% B to wash the column. For APMS experiments, the total MS acquisition time was 60 min, during which mobile phase buffer B (0.1% formic acid in 80% acetonitrile) was ramped from 3% to 28% B over 44 min, followed by a ramp to 45% B over a subsequent 5 minutes, and lastly a ramp to 88% B to wash the column.

For AB, PH, and UB experiments data was collected on an Orbitrap Fusion Tribrid Mass Spectrometer (Thermo Fisher Scientific). The ion transfer tube was set to 300°C, the spray voltage was 1800-2000V, MS1 scan was collected in the orbtrap in profile mode with 120K resolution over a scan range of 400-1600 m/z, a 100 ms maximum injection time, 2e6 AGC target, S-Lens RF of 60. Charges states 2-7 were considered for MS2 scan selection using MIPS filtering, exclusion of undetermined charge states, a 5e3 minimum signal intensity, and a 20s dynamic exclusion with a +/− 10 ppm window. MS2 scans were performed over the course of a 3s cycle, a 1.6 m/z quadrupole isolation, 29% (UB) or 30% (PH and AB) normalized collision energy, HCD fragmentation, a 110 m/z first mass, ion trap detection with a rapid scan rate, 35ms maximum injection time, 1e4 AGC target, centroid data type, and the allowance to injection ions for all available parallelizable time. For APMS experiments data was collected on an Orbtrap Fusion Lumos Tribrid Mass Spectrometer (Thermo Fisher Scientific). The ion transfer tube was set to 275°C, the spray voltage was 2000V, MS1 scan was collected in the orbitrap in profile mode with 240K resolution over a scan range of 350-1350 m/z, a 50 ms maximum injection time, 1e6 AGC target, S-Lens RF of 30. Charges states 2-5 were considered for MS2 scan selection using MIPS filtering, advance peak determination, exclusion of undetermined charge states, a 5e3 minimum signal intensity, and a 20s dynamic exclusion with a +/− 10 ppm window in which 2 occurrences were allowed. MS2 scans were performed over the course of a 1s cycle, a 0.7 m/z quadrupole isolation, 32% normalized collision energy, HCD fragmentation, a 200-1200 m/z scan range, ion trap detection with a rapid scan rate, 18ms maximum injection time, 3e4 AGC target, centroid data type.

#### Proteomics data analysis

All data were searched against the *Mus musculus* UniProt database canonical sequences (taxonomy ID 10090) downloaded September 2^nd^, 2021, and containing 17,085 sequences. Additionally, the human apoE3 and human apoE4 sequences were included. Additionally, APMS data was also searched against the Raph1 mouse sequence. Peptide and protein identification searches, as well as label-free quantitation were performed using the MaxQuant data analysis algorithm (version 2.0.3.0) (35), and all peptide, protein, and modification site identifications were filtered to a 1% false-discovery rate. Default MaxQuant settings were used, with the exception that match between runs was enabled with a window of 2 min (PH and AB), 1 min (UB), or 0.7 min (APMS) for each data type respectively. Searches were performed with a precursor tolerance of 4.5 ppm and a fragment ion tolerance of 0.5 m/z, with trypsin/P specificity (cleaves after lysine and arginine also if a proline follows) allowing for up to 2 missed cleavages. For all datasets a fixed modification of carbamidomethyl cysteine (57.02146 Da), a variable modification of methionine oxidation (15.99491 Da), and a variable modification protein N-term acetylation (42.01056 Da) were used. For phosphorylation experiments an additional variable modification of phosphorylation on serine/threonine/tyrosine (79.96633 Da) was used, while for ubiquitylation experiments an additional variable modification of di-glycine remnant on lysine (114.04293 Da) was applied. Mass spectrometry data files (raw and search results) and annotated spectra have been deposited to the ProteomeXchange Consortium (http://proteomecentral.proteomexchange.org) via the PRIDE partner repository with dataset identifier PXD034346 (user name, password: 8g9e29n4) (36).

Label-free quantification and statistical analysis for both global proteomics and APMS data was performed using MSstats (34). Note, for consistent statistical testing across global proteomics data types, the previously published protein abundance analysis (24) was re-analyzed using a more recent version of the MSstats R-package (version 3.3.10), such that this same MSstats version was used for all global proteomics data types reported here. Enrichment analysis was performed using the enricher function from R packages clusterProfiler (version 3.99.0) (37). The gene ontology terms and annotations were obtained from the R annotation package org.Mm.eg.db (version 3.12.0). We selected the most significant GO term from among redundant GO terms, which were identified using a hierarchical tree based on distances (1-Jaccard Similarity Coefficients of shared genes in GO) between the significant terms, and then cut the tree into clusters containing redundant terms (R functions hclust and cutree, cut height = 0.99). For tissue specificity, proteins with Log2FC(apoE4/apoE3) < −1, and adjusted p-value < 0.05 were input into the “multiple proteins” function in STRING (38). For kinase activity analysis, kinase activities were estimated using known kinase-substrate relationships downloaded from the enzyme-substrate table of OmniPath (39) using the R package OmniPathR[2.0.0] on [01.13.2023]. Kinase-specific data was selected by requiring the modification type to match “phosphorylation” and the gene symbol to be annotated to Gene Ontology term “GO:0016301 kinase activity” in the R annotation package org.Hs.eg.db [3.12.0].

For APMS experiments, interacting protein abundance values derived from MSstats (34) were visualized using Morpheus (https://software.broadinstitute.org/morpheus) and VASP differential protein-protein interaction networks were visualized in Cytoscape (40) with edges between interacting proteins being supplemented from STRING (38).

#### siRNA knockdowns

Pre-designed silencer select siRNAs against APOE, VASP and negative control were ordered from ThermoFisher (Cat. #160728, Cat. #AM16708, Cat. #AM461, respectively). ApoE3 and apoE4 Neuro-2a cells were transfected with siRNA using Lipofectamine RNAiMax (Cat#13778075, Thermo Scientific). Cells were seeded into 6-well plates and transfection protocol was followed according to manufacturer’s instructions (Cat. #13778075, Invitrogen).

#### Constructs

Human GFP-Vasp and mCherry-VASP was gifted by D. Mullins, University of California, San Francisco. Phospho-null mutation VASP S239A was generated using the QuikChange II Site-Directed Mutagenesis Kit (Cat. #200523, Agilent). Primers were designed using the QuikChange Primer Design Program and obtained from Integrated DNA Technologies IDT. VASP S239A Forward Primer: GGCCTCCTCCTGCTTGGCGACTTTCCTGAGTTTG, VASP S239A Reverse Primer: CAAACTCAGGAAAGTCGCCAAGCAGGAGGAGGCC. GFP-VASP and mCherry-VASP constructs were transfected using Polyjet transfection reagent (Cat. #SL100688, Signagen) following the manufacturer’s protocol.

#### Immunoblotting

Cells were lysed in ice-cold RIPA buffer supplemented with cOmplete mini protease inhibitor cocktail (Cat. #1183615300, Roche) and PhosSTOP (PHOSS-RO, Roche). Protein concentration was measured by a Pierce BCA protein assay kit (Cat. #A329645, Life Technologies) and 20ug protein was loaded on a 4%-12% acrylamide gel (Criterion TGX,Cat.#5671095, Biorad). The samples were separated by sodium dodecyl sulfate polyacrylamide gel electrophoresis (SDS-PAGE) and transferred with PVDF membrane (Trans Blot Turbo, Cat. #1704157, Biorad). Membranes are blocked in %5 Milk with Tris-buffered saline with 0.1% Tween 20 detergent (TBST). Primary antibodies were prepared in 5%BSA-TBST and incubated overnight at 4°C. The membranes were washed in TBST and incubated 2 h in species-specific horseradish peroxidase-conjugated secondary antibodies (1:3000-1:5000). Membranes were visualized using chemiluminescence substrate (Pierce ECL, Cat. #32209,Thermo Scientific) and imaged with ChemiDOC MP (Bio-Rad). Antibodies: VASP S239 1:1000 (ab194747, Abcam), VASP 1:1000 (13472-1-AP, Proteintech) ApoE 1:2000 (178479, EMD Milipore), Actin 1:1000 (4970L, CST), Akt 1:1000 (9272S, CST), p-AKT T308 1:1000 (13038S, CST), p-AKT S473 1:1000 (4060S, CST).

#### GFP-VASP Affinity Purification MS

ApoE4 Neuro2a cells were seeded into a 15cm dish. On the day of transfection, cells were transfected with Polyjet using 9ug of GFP-VASP WT and GFP-VASP S239A constructs according to the manufacturer’s instructions (Cat. # SL100688, Signagen). After 8h of transfection, cells were treated either with DMSO control or 15μM H89 and cells were collected after 24h of treatment.

Frozen cell pellets were thawed at room temperature and suspended in 0.5 mL Lysis Buffer [IP Buffer (50 mM Tris-HCl, pH 7.4 at 4°C, 150 mM NaCl, 1 mM EDTA) supplemented with 0.5% Nonidet P 40 Substitute (NP40; Fluka Analytical) and cOmplete mini EDTA-free protease and PhosSTOP phosphatase inhibitor cocktails (Roche)]. Cell suspensions were immediately subjected to one freeze-thaw cycle (at least 20 min on dry ice and partial thawing at 37°C) then refrozen and kept at −80°C until immunoprecipitation. Lysis was completed by partially thawing frozen samples at 37°C and incubating on a tube rotator at 4°C for 30 min. After pelleting debris at 13,000 xg, 4°C for 15 minutes, 25 μL lysate was reserved before proceeding (below). A ThermoMixer C incubator was used to thaw samples at indicated temperatures with constant shaking at 1,100 rpm for 1-2 min; otherwise, samples were kept cold.

For each equilibration/wash step (below), GFP-Trap magnetic agarose beads (20 μL in first plate only; Cat. #GTMA-10, Chromotek) and/or up to 1.0 mL buffer were pre-aliquoted into 96 Deep-well plates, while 0.45 mL lysate was arrayed for each sample. All plates were kept on ice until loaded onto the KingFisher Flex (KFF) Purification System (Thermo Scientific) for automated processing as follows: GFP-Trap beads were equilibrated twice with IP Wash Buffer (IP Buffer supplemented with 0.05% NP40) and incubated with cell lysates for 2 hours. Protein-bound beads were washed three times with IP Wash Buffer and then once with IP Buffer before elution. After transferring to 1.5-mL lobind tubes, beads were suspended with 50 μL 0.05% RapiGest in IP Buffer and gently agitated on a vortex mixer at room temperature for 30 min to elute proteins and briefly rinsed in 25 μL IP Buffer to recover residual proteins; both were combined for further processing (below). The KFF is operated in a cold room to maintain a 4°C temperature during immunoprecipitation. Automated protocol steps were performed using the slow mix speed and the following mix times: 30 seconds for equilibration/wash steps, 2 hours for binding, and 1 minute for final bead release. Three 10 second bead collection times were used before bead transfer to the next plate.

Using a 96 well PCR plate, samples were incubated at 37°C in a thermal cycler or at room temperature in a MixMate incubator with shaking at 1,100 rpm for the following steps: Denaturation and reduction at 37°C for 30 minutes with 2 M urea and 1 mM DTT in 50 mM Tris-HCl pH 8.0, alkylation at room temperature for 45 minutes with 3 mM iodoacetamide and quenching for 10 minutes with 3 mM DTT. Trypsin (0.5 μg/μL; Promega) was added twice (1.0 μL and 0.5 μL) and further incubated at 37°C for 4 hours and then another 2 hours. Peptides were acidified with trifluoroacetic acid (TFA, 0.5% final, pH < 2.0) and desalted at room temperature using a BioPureSPE Mini 96-well plate (20mg PROTO 300 C18; The Nest Group). Briefly, desalting columns were sequentially equilibrated with 100% methanol; 80% acetonitrile (ACN), 0.1% TFA and 2% ACN, 0.1% TFA before passing acidified samples through columns twice and subsequently washed with 2% ACN, 0.1% TFA and 0.1% formic acid (FA). 0.2 - 0.3 mL of buffer was added at each step. Peptides were eluted twice with 50% ACN, 0.1% FA (0.12 mL total, divided) and dried under vacuum centrifugation (CentriVap Concentrator, Labconco). The desalting plate was centrifuged at 2,000 rpm for 2-3 minutes. All buffers were prepared with HPLC or LC-MS grade reagents.

#### Immunofluorescence

Neuro2a cells were cultured on coverslips. After one PBS wash, cells were fixed in 3.7% paraformaldehyde for 10 min. Then, cells were permeabilized with 0.5% TritonX-100 for 5 min, followed by the addition of 100nM Phalloidin to the cells. Phalloidin staining protocol was followed according to manufacturer’s instructions (Cat. #PHDH1, Cytoskeleton Inc). For VASP staining, cells were blocked in 5% normal goat serum (NGS) for 15min at room temperature. VASP antibody (13472-1-AP, Proteintech) was prepared in 5% NGS (diluted in 1:200) and incubated overnight at 4°C. The next day, cells were washed with PBS and incubated with the secondary antibody (diluted in 1:500, 5% NGS) at room temperature for 45min. The coverslips were mounted on glass slides using DAPI Fluoromount-G (Cat. #0100-20, Southernbiotech) and imaged using a Visitech iSIM superresolution module and Hamamatsu Quest camera on a Nikon Eclipse Ti-E with a 100x 1.45NA objective. Images were deconvolved using Microvolution and neurite length was calculated using NeuronJ plugin in Fiji, as previously described (41). NeuronJ traces the neurite by finding two points, one at the start and one at the end and calculating the optimal intensity path. Neurite lengths were quantified using Prism graphpad with three independent biological replicates.

#### Conditioned media experiment

Neuro2a-apoE3 and Neuro2-apoE4 cells were seeded in a T75 flask and after 72h, media containing either apoE3 or apoE4 were collected from the T75 flask and briefly centrifuged to remove debris. Next, Neuro2a-apoE3 and Neuro2a-apoE4 cells were seeded into 6-well plates with a 50% density, and to the cells was added either normal media, or conditioned media (from apoE3 or apoE4 cells). Cells were incubated with the respective media for three days, then the cells were lysed, protein concentration was measured with BCA assay, and p-VASP levels were measured by immunoblotting.

#### Primary neuron lysates

Primary neuron cultures were prepared from embryonic day 18–20 or postnatal day 0 (P0) pups of three genotypes: 1. Neuron-specific enolase promoter-driven APOE4-expressing (NSE-APOE4) transgenic mice, 2. Neuron-specific enolase promoter-driven APOE3-expressing (NSE-APOE3) transgenic mice, and 3. Wild type C57BL/6 mice. 48 well culture plates (Corning 3548) were coated with Poly-L-Lysine (Sigma P4707) overnight and washed four times with water. The cortex and hippocampus were isolated, and the dissociated cells were plated at a density of 315,000 cells/cm 2 (300,000 cells/ well) in Neurobasal medium (Gibco 21103-049) supplemented with B27 (Gibco 17504-044), 100 U/ml penicillin G, 100 μg/ml streptomycin (Gibco 15140-122), and 1% Glumax (Gibco 35050-061). Primary neurons were cultured in vitro for 7 days before treatment with 0.1% DMSO three times during the following week. After 14 days primary neurons were collected by adding 80μl/ well of Low Detergent Buffer [50mM Tris, pH7.5, 150mM NaCl, 1% NP40, 0.5% Sodium Deoxycholate, 0.1% SDS, 1 pellet of Protease Inhibitor (Roche 11836145001) per 25ml], scratched off the plate and homogenized with a P200 pipet tip. Lysates were transferred to labeled Eppendorf tubes and spun at 14,000rpm for 5min at room temperature. The neuron lysates were collected, stored at −20°C and analyzed in Western blot assays. Due to limited protein amounts lysates from different cultures were randomly pooled to provide sufficient protein amounts. This resulted in a single lysate for apoE3 neurons, and two lysates for apoE4 neurons.

#### Statistical analysis

For non-proteomics data a two-tailed t-test was performed when two groups were compared and one-way ANOVA or two-way ANOVA with repeated measures was performed when multiple groups were compared, according to different conditions. Where ANOVA-tests revealed significant differences among means, a *post-hoc* Sidak test was performed to identify significant differences. Statistical analyses were performed with Prism software (GraphPad Software, Inc.). Differences were considered significant at *p < 0.05, **p < 0.01 and ***p < 0.001 ****p < 0.0001. SuperPlots in Figure 5 were generated in GraphPad Prism (42).

## Results

### Post-translational modification networks are highly regulated by apoE4

Our previous work has shown that apoE allele status can significantly regulate the proteome, resulting in mitochondrial dysfunction and altered cellular metabolism (24). To further define the proteomic impact of apoE4, we analyzed differences for thousands of protein phosphorylation and ubiquitylation sites between isogenic mouse Neuro-2a cells expressing human apoE3 or apoE4 (**Fig. 1 A-B, Table S1**). These neuron-like cells have been extensively characterized and shown to recapitulate numerous phenotypes observed in AD patients, including increased Amyloid β (Aβ) (10), tau hyperphosphorylation and NFTs (43), and mitochondrial dysfunction (24). Additionally, we found that apoE4 expression in Neuro-2a cells resulted in depletion of proteins normally expressed in the hippocampus (FDR = 4.5×10^−3^), a brain region that incurs neuronal loss in AD patients, further supporting the relevance of these cells as an AD model (**Table S2**).

**Figure 1.**
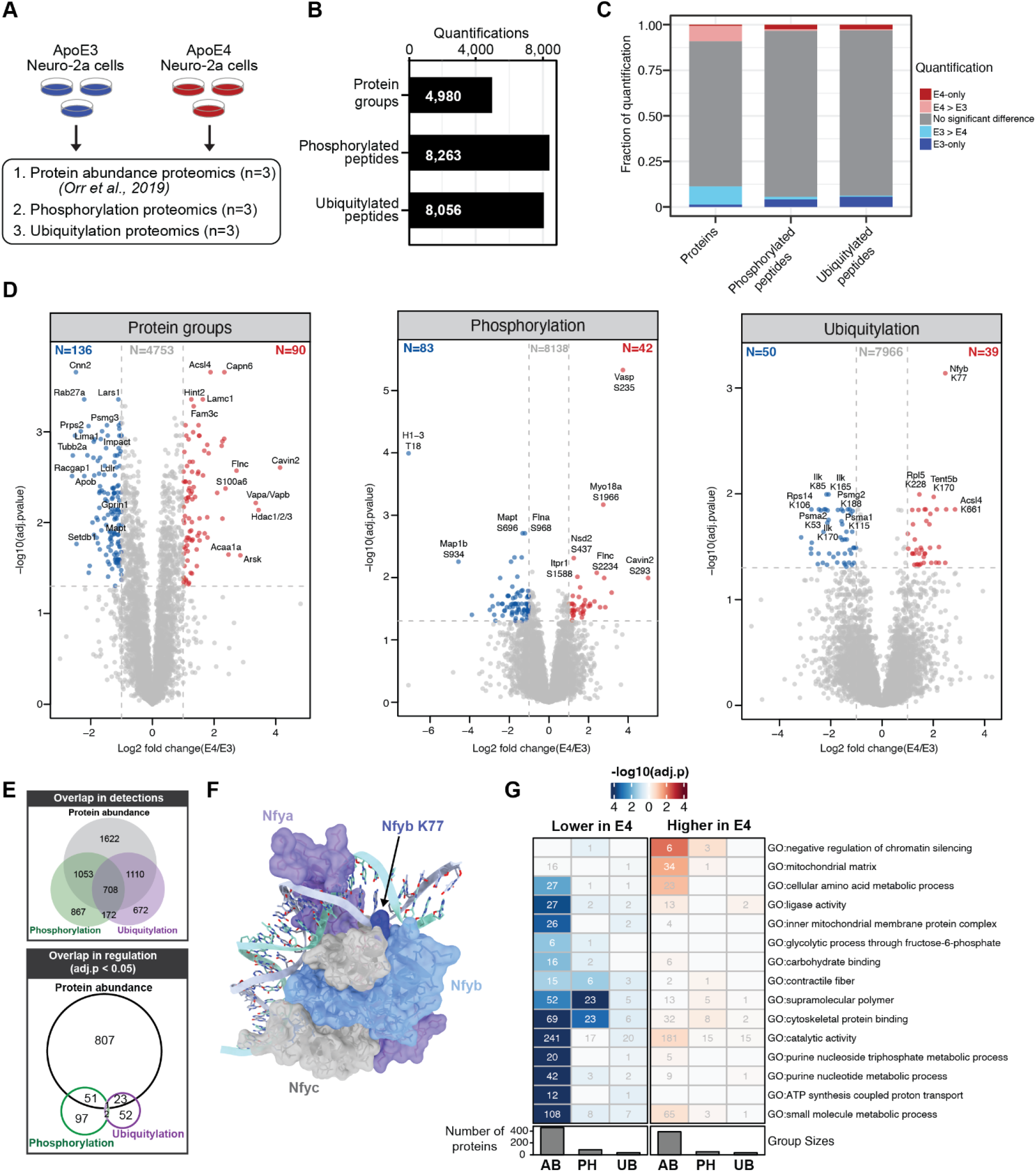
Experimental overview. In all panels, differential regulation is defined as proteins or PTMs with adj.p < 0.05. (**A**) ApoE3 and apoE4 Neuro-2a cells were subjected to proteomic analysis in biological triplicate, measuring protein abundance (data re-analyzed from our previous report (24)), phosphorylated peptides, and ubiquitylated peptides. (**B**) Total number of unique quantifications for each analysis method. (**C**) Fraction of differentially regulated qualifications for each analysis method. When a peptide or protein was only detected in one cell type it was designated as E3-only or E4-only. (**D**) Volcano plots for quantified protein, phosphorylated peptides, and ubiquitylated peptides. Colored data points indicate analytes with adj.p < 0.05 and a log_2_ fold change > |1|. A selection of significantly regulated analytes are annotated with their gene name and the modified amino acid position. (**E**) Overlap of proteins or PTM-containing proteins from all quantified proteins/PTMs (top), or only from those proteins/PTMs that were differentially regulated (adj.p < 0.05) (bottom). (**F**) Structure of the Nfy trimeric protein complex bound to DNA (PDB: 4AWL). Nfyb is shown in light blue, with the position of K77 highlighted in dark blue. Nfya is shown in purple, and Nfyc is shown in grey. (**G**) Gene ontology enrichment analysis. Enrichment analysis was performed on proteins or PTMs with adj.p < 0.05.

While there was a high degree of overlap in the proteins detected across data types, we observed apoE allele regulation to impact highly unique subsets of the proteome across the multiple modes of regulation (**Fig. 1D-E**), as has been observed in the analysis of AD patient brain tissue (44). From the ubiquitylation data, we find several components proteasome to have lower ubiquitylation levels in apoE4 cells (e.g. Psmg2, Psma1, and Psma2). The most significantly regulated ubiquitylation detected was K77 on the nuclear transcription factor Y subuit beta (Nfyb) protein. Nfyb functions as part of a trimeric protein complex, along with subunits alpha and gamma, to bind to CCAAT motifs in the promoter regions of DNA. Interestingly, our previous publication reporting only the protein abundance data found that Nfy was predicted to regulate the differentially expressed proteins observed between apoE3 and apoE4 Neuro-2a cells (24). We evaluated this positioning of this ubiquitylation site within the structure of the trimeric Nfy protein complex (PDB: 4AWL) (45), and find that Nfyb K77 side chain fits within the DNA minor groove, such that ubiquitylation of this residue would likely inhibit binding of this complex to DNA, and consequently its Nfy transcriptional activity.

Gene ontology (GO) analysis of the datasets revealed a modest set of enriched terms, with cytoskeletal protein binding being among the most highly enriched terms among both protein abundance and phosphorylation sites that were found to be lower in apoE4 cells (**Fig. 1G, Table S2**). Indeed, in the case of phosphorylation sites, many of the most significantly regulated sites in our dataset, both higher and lower in apoE4 cells, are associated with cytoskeletal regulation (e.g. Map1b, Mapt, Flna, Flnc, Myo18a, and Vasp).

To better enable biological interpretation of regulated phosphorylation sites, they can be mapped to known upstream kinases. Here we mapped our detected phosphorylation sites from mouse to their orthologous human phosphorylation sites, mapped these sites to kinases, and then predicted kinase activities based on the differential regulation of these phosphorylation sites (**Fig. 2A, Table S2**) (46). Evaluation of kinase activities from our dataset predicted strong inactivation of many components of the MAPK pathway in apoE4 cells, with activation of many kinases (GUCY2C, PRKD2, PRKG2, PRKG1) being driven by the strong increase in VASP S235 phosphorylation. To test the accuracy of these predictions we selected one kinase, AKT2, for which we find 9 substrate phosphorylation sites that predict an increase in AKT2 kinase activity in apoE4 Neuro-2a cells. In support of these predictions, we tested apoE3 and apoE4 lysates with antibodies that recognize different AKT phosphorylation sites and found that AKT phosphorylation was highly increased in apoE4 cells on two sites known to regulate its kinase activity (T308/T309/T305 and S473/S474/S472 on AKT1/2/3, respectively, **Fig. 2B**).

**Figure 2.**
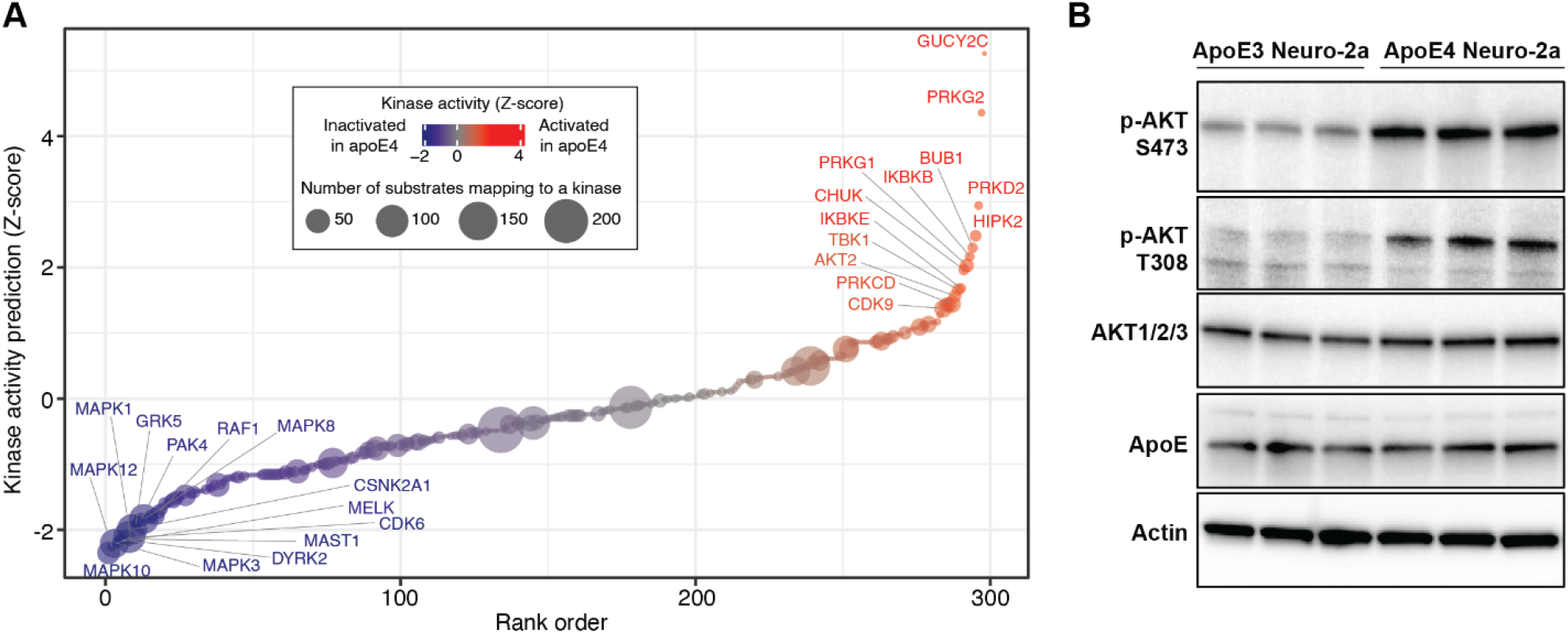
Analysis of kinase activities. (**A**) Kinase activity prediction analysis. Dot size is proportional to the number of substrates detected in the phosphoproteomics experiments, and dot color represents the predicted level of inactivation or activation of each kinase. (**B**) Western blot measuring AKT phosphorylation at S473/S474/S472 and T308/T309/T305 on AKT1/2/3, AKT1/2/3 total protein, and apoE protein levels in Neuro-2a apoE3 or apoE4 cell lysates. Actin serves as a loading control.

### The cytoskeletal protein VASP is phosphorylated at S235 in an apoE4-dependent manner

We hypothesized that the enrichment of cytoskeletal proteins in both the protein abundance and phosphorylation analyses may provide mechanistic insights into the established cytoskeletal defects previously observed in apoE4 cells. Intriguingly, the most significantly regulated phosphorylation site in our data was Vasodilator-stimulated phosphoprotein (VASP) S235, which was 14-fold higher in apoE4 cells (**Fig. 1D**), and has previously been detected in apoE4-carrier AD patient brains (47).

VASP is a member of the conserved ENA/VASP family of proteins regulating diverse cellular processes such as cell adhesion, neurite initiation, axon outgrowth and actin cytoskeletal dynamics (48). VASP is composed of three conserved domains: an N-terminal Ena/VASP homology-1 (EVH-1) domain, a central poly-proline rich domain (PPR) and a C-terminal EVH-2 domain (49, 50) (**Fig. 3A**). While EVH-1 interacts with proteins such as zyxin and vinculin, which are important for cytoskeletal regulation, the PRR domain mediates the interaction with profilin-actin complexes to enhance actin polymerization (50, 51). Beyond VASP itself, we found that many proteins related to VASP biology were also differentially regulated phosphorylation in apoE4-expressing cells, including zyxin (Zyx), drebrin (Dbn1), actin-binding LIM1 protein (Ablim1) and Cofilin-1 (Cfn1) proteins (**Fig. 3B**). Many of these had previously been linked to AD pathogenesis or nervous system development, including Dbn1 (52), Cfn1 (53), and Zyx (54–57). In line with this, our global proteomic analysis showed that abundance levels of these four proteins are reduced in Neuro-2a apoE4 cells (**Fig. 3B**).

**Figure 3.**
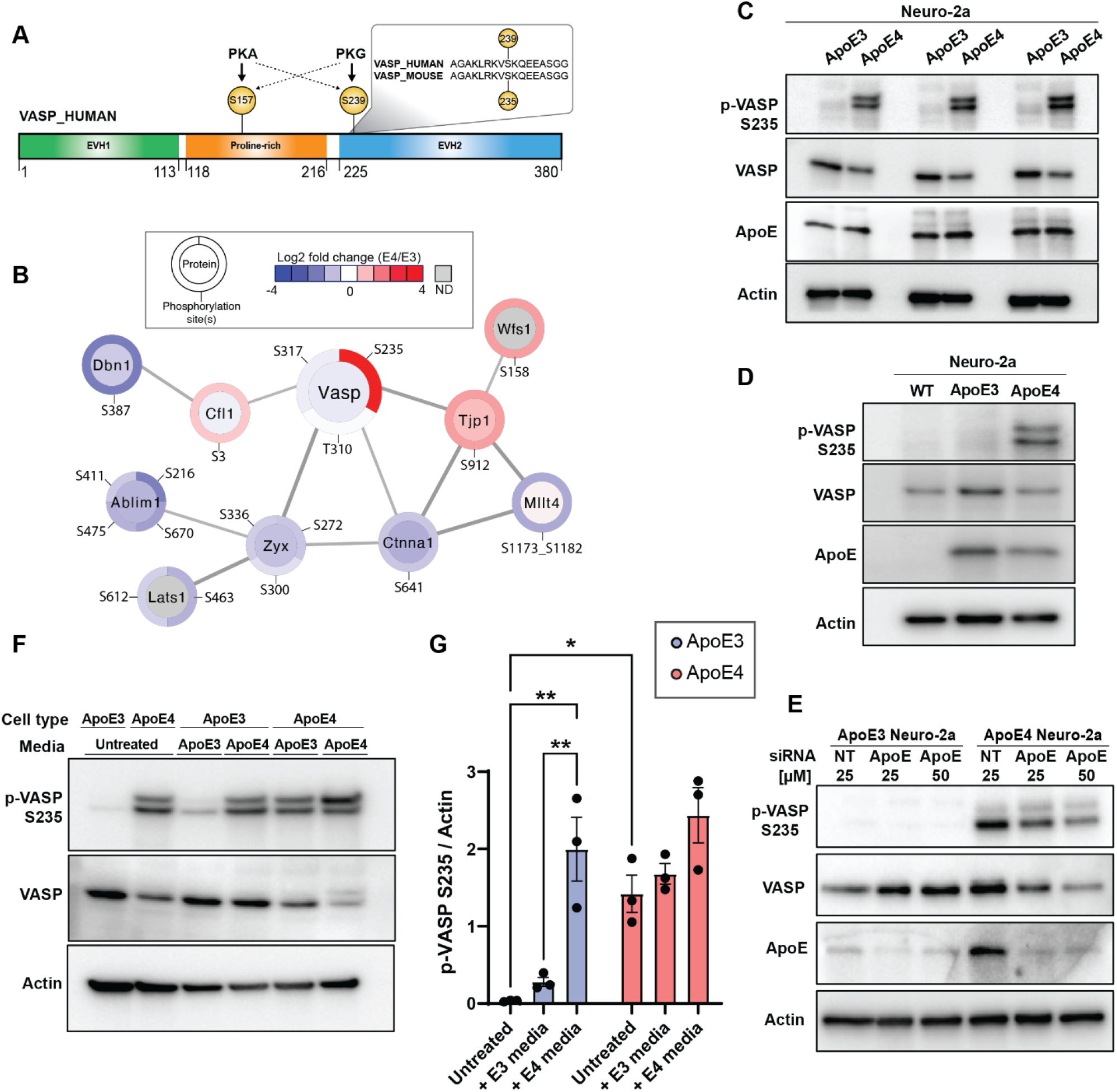
VASP is phosphorylated at S235 in an apoE4-dependent manner. (**A**) VASP protein domain organization: EVH1: Ens/VASP Homology 1, Poly-Proline rich domain (PPR), EVH2: Ena/VASP homology 2. VASP proteins are the substrates of PKA or PKG. VASP S235 (mouse) and VASP S239 (human) are conserved across species. (**B**) Quantification of selected phosphorylation sites and protein abundance differences for proteins known to interact with VASP based on the STRING database. Protein abundance quantification is displayed in the interior circle. Phosphorylation site amino acid position and quantification annotations are displayed on the border, ND:not detected. (**C**) Phosphorylation of VASP at S235 in apoE4 Neuro-2a cells validation by immunoblotting (n=3 biological replicates), with actin as a loading control. (**D**) VASP phosphorylation at S235 in Neuro-2a cells in the absence of apoE (WT) or different apoE alleles. Actin serves as a loading control. (**E**) Western blot measuring VASP S235 phosphorylation, VASP protein and apoE protein upon knockdown of apoE with the increasing concentrations of siRNA in apoE3 Neuro-2a and apoE4 Neuro-2a cells. NT indicates a non-targeting siRNA as a negative control. Actin serves as a loading control. (**F**) Western blot of VASP S235 phosphorylation and VASP total protein in Neuro-2a apoE3 or apoE4 cells cultured with normal media (untreated) or with conditioned media from Neuro-2a apoE3 or apoE4 cells. Actin serves as a loading control. (**G**) Quantification of VASP S235 phosphorylation relative to actin for each sample. Data are mean ± s.d. from n=3 biological replicates per group * p<0.05; ** p<0.01.

We validated the increase in VASP S235 phosphorylation in apoE4 cells by immunoblotting, and found that it occurred without a concomitant increase in VASP protein abundance (**Fig. 3C**). This increase in phosphorylation was unique to apoE4 expressing cells, as VASP S235 phosphorylation was not detectable in wild-type (WT) Neuro-2a cells that lack exogenous human apoE expression (**Fig. 3D**). Furthermore, to evaluate if this phosphorylation site is present in apoE4 cells beyond Neuro-2a, we measured VASP S235 phosphorylation from primary neurons derived from either WT mice, or knock-in mice with neuron-specific expression of either human apoE3 or apoE4. Here we observed this phosphorylation site in two pools of apoE4 primary neuron lysates, as well as to a lower extent in WT mouse lysates (**Fig. S1**).

To further confirm the role of apoE4 in promoting VASP S235 phosphorylation, we performed siRNA knockdown of apoE and observed up to a 50% reduction in this phosphorylation site in apoE4 cells (**Fig. 3E, S2**). Lastly, while apoE can escape the secretory pathway to modulate intracellular biology, it is predominantly a secreted protein. Thus, we evaluated if conditioned media, which contains both apoE and a variety of other secreted factors, can regulate VASP S235 phosphorylation. ApoE3 or apoE4 Neuro-2a cells were cultured for 3 days using either normal media, or conditioned media from either apo3 or apoE4 Neuro-2a cells. We find that conditioned media from apoE4 Neuro-2a cells significantly increases VASP S235 phosphorylation levels in apoE3 cells (**Fig 3F-G**). These results suggest that a secreted factor from apoE4 cells (potentially apoE itself) is responsible for initiating the cellular signaling cascade that induces VASP S235 phosphorylation.

### VASP S235 phosphorylation is mediated by PKA

Phosphorylation of VASP at serine 235 (mouse), or the orthologous serine 239 (human), is known to impair F-actin accumulation, cell adhesion, and filopodia formation (58) (**Fig. 3A**). The serine threonine kinases PKG and PKA can phosphorylate VASP at S235. PKG is the preferred kinase for this site (59) and consequently was predicted to be activated in apoE4 cells (**Fig. 2A**), while PKA has been shown to preferentially phosphorylate VASP S157 (**Fig. 3A**) (48). In order to understand which protein kinase is responsible for VASP S235 phosphorylation in apoE4 Neuro-2a cells, we first treated cells with two different PGK inhibitors RRKAARE (**Fig. 4A**) or KT5823 (**Fig. S3**); however no decrease in phosphorylation was observed. In contrast, treatment of cells with two different PKA inhibitors, H89 (**Fig. 4B-C**) or KT5720 (**Fig. S3**), both dramatically reduced VASP S235 phosphorylation. These results suggest that PKA is responsible for the majority of VASP S235 phosphorylation and is the preferential kinase for this site in apoE4 Neuro-2a cells.

**Figure 4.**
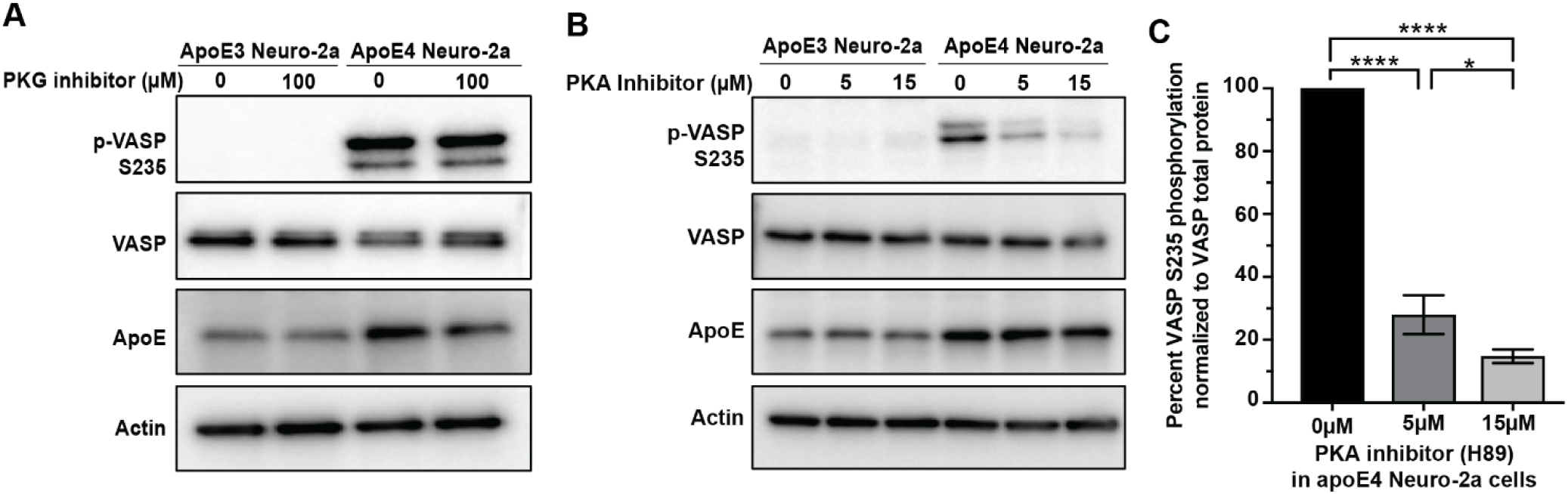
VASP S235 is phosphorylated by PKA. Immunoblots of VASP S235 phosphorylation in apoE3 and apoE4 Neuro-2a cells upon kinase inhibition. (**A**) 24h inhibition with the PKG inhibitor, RKRARKE. (**B**) 24h inhibition with increasing concentrations of the PKA inhibitor H89. Actin serves as a loading control. (**C**) Quantification of VASP S235 phosphorylation levels from panel B. Data are mean ± s.d. from n=3 biological replicates per group * p<0.05; **** p<0.0001.

### VASP interactions with actin cytoskeletal proteins are regulated by PKA

To investigate the role of VASP S235 phosphorylation in apoE4 Neuro-2a cells, we performed affinity purification-mass spectrometry (APMS) to identify how VASP protein-protein interactions (PPIs) are regulated by its phosphorylation status. We expressed wild-type human VASP (hVASP WT), which was highly phosphorylated at the orthologous S239 position in apoE4 Neuro-2a cells (**Fig. S4**), similar to what we observed for the endogenous mouse VASP S235 phosphorylation. Separately, we expressed human VASP with an S239A mutation (hVASP S239A) which prohibits phosphorylation at this position. APMS experiments were performed in the presence or absence of PKA inhibition by H89 (**Fig. 5A**).

**Figure 5.**
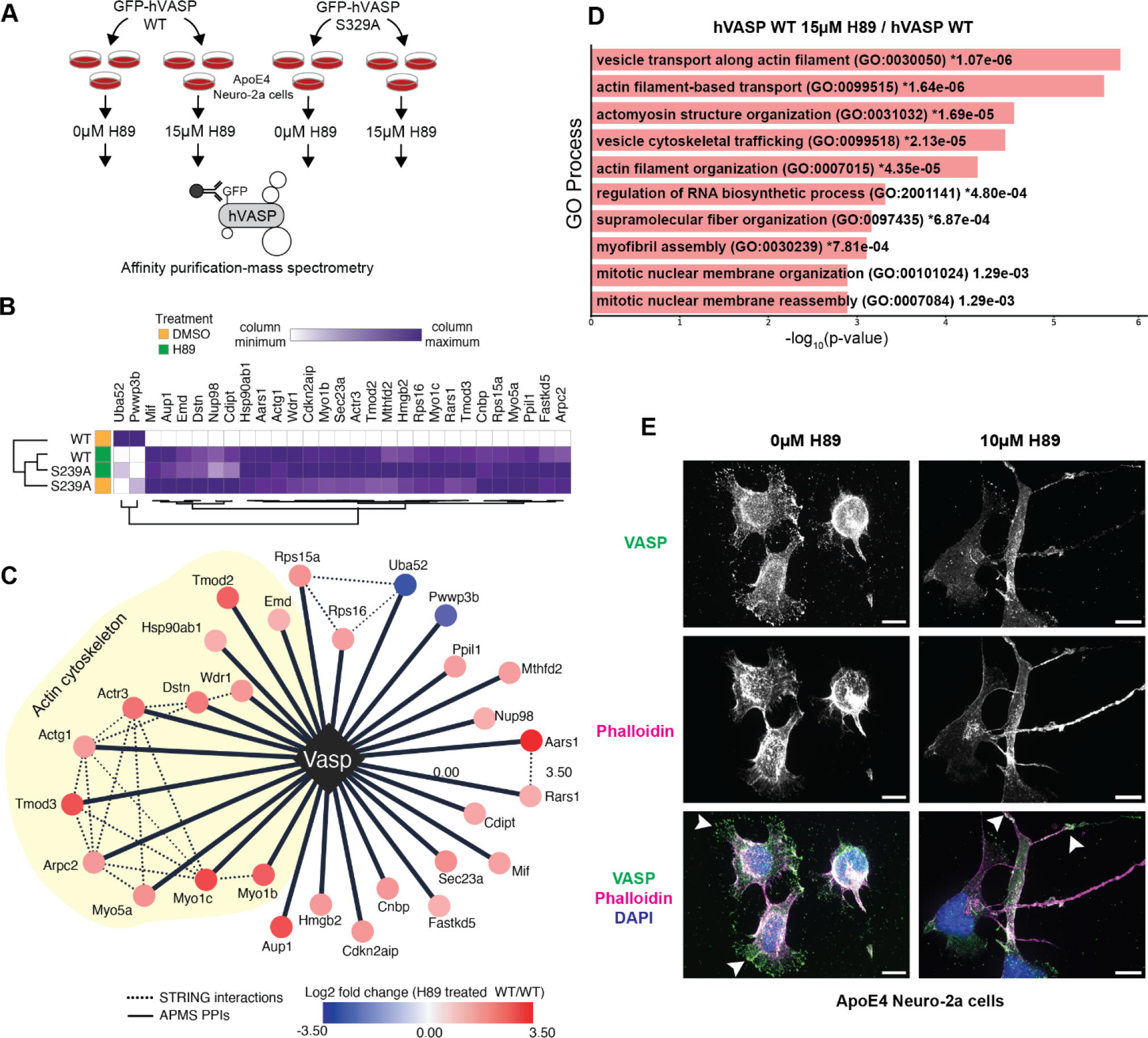
Phosphorylated VASP and phospho-null VASP protein interacting networks. (**A**) GFP-tagged human VASP (hVASP) WT and hVASP S239A are expressed in apoE4 Neuro-2a cells in triplicate in the absence or presence of 15μM H89. GFP is used as a tag to pull down VASP protein interacting partners which were identified using MS. (**B**) Heat map represents scaled protein abundance values for a selection of interacting proteins specifically regulated by hVASP S239 phosphorylation. (**C**) Comparison of hVASP WT with H89 treated hVASP WT protein interaction network for proteins displayed from panel B. (Red, increase; blue, decrease). (**D**) Bar plot of Gene Ontology analysis of proteins differentially purified from hVASP expressing apoE4 Neuro-2a cells in the absence or presence of H89. Proteins in Gene ontology analysis using EnrichR were determined by a p < 0.05 and a log_2_ fold change >1 and sorted with p-value ranking. (**E**) Localization of VASP in apoE4 Neuro-2a cells in the absence or presence of 10μM H89 for 24h. Phalloidin staining is for F-actin. DAPI: Nuclear staining. The scale bar is 10μm.

To prioritize PPIs dependence on hVASP S239 phosphorylation status, we filtered the APMS data for proteins with differential binding (>2-fold) between WT hVASP and S239A hVASP, as well as differential binding upon dephosphorylation of WT S239 by H89 treatment. Additionally, to control for H89 effects on hVASP PPIs that may not be related to its S239 phosphorylation status, we further filtered PPIs to those that were relatively unchanged (<2-fold) in S239A mutant cells upon H89 treatment. These criteria resulted in a set of 29 proteins whose interactions were consistent across all conditions except when hVASP was in a highly phosphorylated state at position S239 (i.e., WT hVASP) (**Fig. 5B, Table S3**). Blocking hVASP S239 phosphorylation increased PPIs with components of the actin cytoskeleton (**Fig. 5C**). For instance, hVASP interactions with microfilament motor activity proteins Myo1b, Myo1c, and Myo5a were increased following PKA inhibition in apoE4 Neuro-2a cells. Myo1b is involved in neurite outgrowth, cell migration, axon formation, and growth cone dynamics (60). Actin nucleating and branching protein Arp2/3 and its component Actr3, as well as Actin protein Actg1, also preferentially interacted with hVASP upon S239 dephosphorylation by H89 (**Fig. 5C**). In line with this, GO analysis of changes in this protein interaction network showed an enrichment of pathways associated with actin filament transport, trafficking, and reorganization (**Fig. 5D**).

We next asked whether the observed changes in PPIs were influenced by VASP subcellular localization in apoE4 Neuro-2a cells, comparing VASP and F-actin subcellular localization patterns in apoE4 Neuro-2a cells in the absence or presence of H89 (**Fig. 5E**). When VASP S235 was phosphorylated, it was predominantly localized at the leading edge and in the cytosol (**Fig 5E**). However, upon dephosphorylation of S235 by H89 treatment, VASP became less cytosolic and was enriched as clusters in F-actin containing neurites (**Fig. 5E**). Taken together with APMS data, these findings suggest VASP phosphorylation at S235 by PKA alters actin cytoskeleton protein interactions and subsequently VASP subcellular localization in apoE4 Neuro-2a cells.

### Inhibition of PKA promotes filopodia formation and neurite outgrowth in apoE4 cells

Previous studies have shown that apoE4 expression inhibits neurite outgrowth (28, 29), a process which is initiated by robust filopodia formation (61). The observation of long F-actin neurite structures upon PKA inhibition in apoE4 Neuro-2a cells led us to hypothesize that VASP S235 phosphorylation could be an apoE allele-specific mechanism regulating neurite outgrowth and synaptic remodeling through actin cytoskeleton-dependent processes. To test this, we used H89 to inhibit VASP S235 phosphorylation in both apoE3 and apoE4 Neuro-2a cells, and quantified the impact on filopodia formation and neurite outgrowth. As expected, under normal conditions, apoE4 cells had moderately shorter filopodia than apoE3 cells. However, PKA inhibition resulted in a dramatic increase in the length of filopodia in apoE4 cells, some of which appeared to form neurites (**Fig. 6A,C**). Next, we measured neurite outgrowth under these same conditions and observed that indeed, neurite outgrowth was significantly increased in H89-treated apoE4 cells as compared to control apoE4 cells (**Fig. 6B,D**). In contrast, H89 treatment cells did not significantly increase neurite outgrowth in apoE3 cells. Taken together, these results show that in apoE4 cells, depletion of VASP S235 phosphorylation by inhibition of PKA leads to extended filopodia length and subsequently neurite outgrowth.

**Figure 6.**
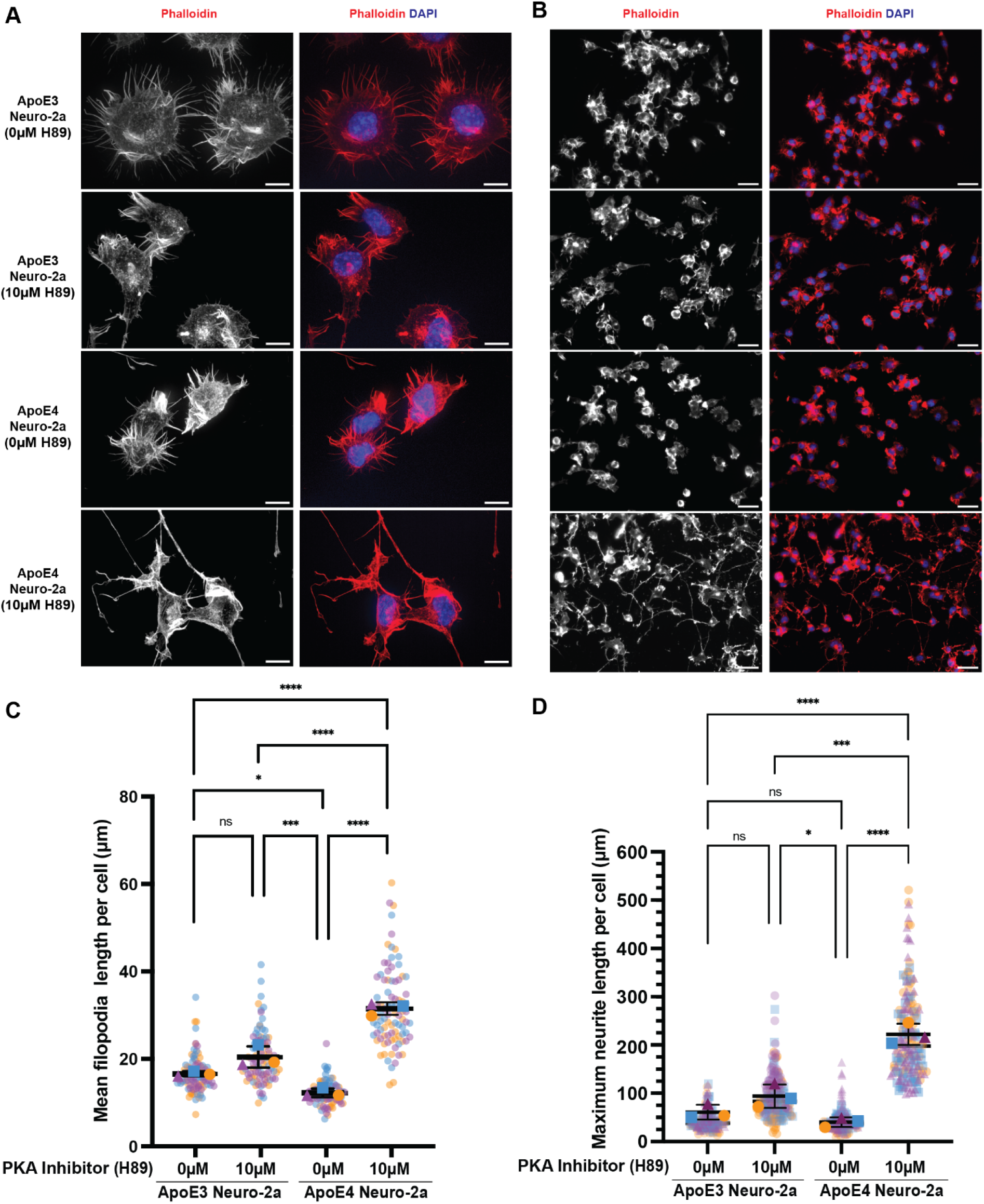
PKA inhibition increases filopodia and neurite length. (**A-B**) Phalloidin staining of apoE3 or apoE4 expressing Neuro-2a cells in the absence or presence of 10μM PKA inhibitor (H89), DAPI: Nuclear staining. Images in panel B are lower magnification to show more cells; scale bar is 10μm (A) and 200μm (B). (**C-D**) Measurement of mean of filopodia and max neurite length per cell using NeuronJ (41). Data represents n=3 independent samples (biological replicates). Each small data point (purple, blue, yellow) represents the length per cell, color-coded by the sample; error bars are ±s.d. of the biological replicates. The triangle represents the average of purple data points, the square represents the average of blue data points, the sphere represents the average of yellow data points. *P* values were calculated using the means of the biological replicates (ns: non-significant, * p<0.05; *** p<0.001; **** p<0.0001; with one-way ANOVA analysis with Tukey’s multiple comparison test).

## Discussion

Neurons can undergo dramatic morphological changes, forming abundant neurite extensions, axonal elongations, synaptic spines and synapses, all guided by cytoskeletal elements. Disruptions to these cytoskeletal processes are a common feature of AD neuropathogenesis, leading to abnormal actin-containing inclusions, NFT formation and synaptic deficits (62). These phenotypes can be specifically regulated by apoE4 expression, which has been documented to result in cytoskeletal impairments via mechanisms that have remained unclear (27–31). The global phosphoproteomics analysis presented here provides insights into the molecular mechanisms of this process, implicating PKA and the actin-interacting protein VASP.

Our findings demonstrate that apoE4 expression results in differential phosphorylation resulting in rewiring of the phosphorylation signaling architecture. Our observations in Neuro-2a cells are supported by clinical data, in which phosphorylation signaling was found to readily distinguish AD patients from other conditions, including progressive supranuclear palsy (PSP), mild cognitive impairment, or individuals with high pathology but low cognitive impact (44). Additionally, several phosphoproteomics studies have demonstrated that PKA activity is up-regulated in AD patient brains (44, 47). Human VASP phosphorylation at S239 (the orthologous site to mouse S235) has also been detected in a handful of patients carrying the apoE4 allele (47), however likely due to the high density of lysine and arginine residues proximal to this phosphorylation site, insufficient detections of this site have been reported in clinical cohorts. Thus, further studies utilizing alternative proteolytic enzymes or detection methods would be required to statistically correlate this phosphorylation site with apoE4 allele status, AD diagnosis, or cytoskeletal dysfunction in patients.

Previous work has established a role for apoE4 in promoting PKA activity. For example, in smooth muscle cells and mouse embryonic fibroblasts, extracellular apoE4 can bind to LRP1, resulting in increased cAMP levels and consequently activation of PKA (63). Internalization of apoE4 through the LRP receptor has been shown to reduce neurite extension and branching in Neuro-2a cells (27). Furthermore, incubation of rat primary hippocampal neurons with extracellular apoE4 results in the stimulation of the cAMP response element-binding protein (CREB) (64), a downstream PKA target. Our results build upon these observations, and indicate that of the many potential downstream PKA targets, VASP S235 phosphorylation is dramatically upregulated in the apoE4-expressing cells. Interestingly, we also find that factors present in apoE4 conditioned media are sufficient to drive VASP S235 phosphorylation, however it is unclear if this is regulated by secreted apoE4, or other factors present uniquely in the apoE4 media.

We found that PKA inhibition increased the interaction of VASP with proteins such as Myo1b, Myo5a, Tmod2 and Tmod3, which are involved in actin filament organization, vesicle transport along the actin filaments and actomyosin structure organization. This suggests that apoE4-dependent cytoskeletal dysfunction through VASP phosphorylation at S235 could be reversed by PKA inhibition. There is evidence that in response to an increase in cAMP levels in neurons, PKA subunit C translocates from the cytosol to the plasma membrane to phosphorylate its substrates (65). As VASP is localized at the leading edge, phosphorylation of S235 by PKA may prevent VASP from being involved in filopodia formation and neurite initiation.

Filopodia are composed of VASP clusters, parallel filament actin bundles, and other actin binding proteins such as lamellipodin (66). Together with microtubules, filopodia are required for neurite initiation and axon extension, which are essential processes for neuronal migration, synapse formation and axon outgrowth and guidance (66–69). Accordingly, our abundance and phosphoproteomics data suggest that numerous proteins involved in actin cytoskeleton organization, actin filament-based processes, cell leading edge, postsynaptic actin cytoskeleton organization and microtubule-based transportation are reduced in apoE4 Neuro-2a cells. In contrast, proteins involved in actin filament fragmentation, actin filament severing and depolymerization processes are elevated in apoE4 Neuro-2a cells. Surprisingly, in contrast to our findings, previous studies have shown that neurite initiation and length are reduced upon PKA inhibition in zebrafish neurons and rat hippocampal neurons (70, 71), however these studies did not evaluate the impact of apoE4 on these phenotypes.

In summary, our results suggest a mechanism where increased PKA activity and VASP S235 phosphorylation may limit cytoskeletal dynamics in apoE4 neurons, and consequently limit the capacity for such neurons to engage in synaptic remodeling. Importantly, our results also indicate that this phenotype can be reversed upon PKA inhibition. Thus, further elucidation of the role of VASP and PKA in actin cytoskeletal reorganization may provide valuable insights into apoE4-dependent neuropathological mechanisms.

## Abbreviations

PTMs: Post-translational modifications
apoE: Apolipoprotein E
VASP: Vasodilator-stimulated phosphoprotein Neuro-2a
KD: Knockdown
WT: Wild type
Aβ: Amyloid β
NFT: Neurofibrillary Tangle
PPIs: protein-protein interactions
APMS: Affinity purification-mass spectrometry

## Acknowledgements

We thank Yadong Huang, Kristen Skruber, Anke Meyer-Franke, and Michael McGregor for helpful discussions. We thank Einar Krogsaeter and Kliment Verba for assistance with figures. This work was supported by an NIH grant R01AG059751 to RWM, NJK, and DLS, an NIH grant R35GM118119 to RDM, and funding from the Howard Hughes Medical Institute Investigator Program to RDM.

## Data availability

Mass spectrometry data files (raw and search results) have been deposited to the ProteomeXchange Consortium (http://proteomecentral.proteomexchange.org) via the PRIDE partner repository with dataset identifier PXD034346 (user name, password: 8g9e29n4) (36).

## Author contributions

**Zeynep Cakir**: Conceptualizing, investigation, visualization, validation, formal analysis, writing-original draft, writing - reviewing & editing. **Samuel J. Lord**: Investigation, Visualization, Resources. **David Jimenez-Morales**: Formal analysis. **Gwendolyn M. Jang**: Investigation. **Benjamin J. Polacco**: Formal analysis and visualization. **Manon Eckhardt**: Writing - Review & Editing. **Billy W. Newton**: Investigation. **Yuan Zhou**: Formal analysis and visualization. **Adam L. Orr**: Investigation. **Jeffery R. Johnson**: Methodology, investigation, software, supervision. **Alexandre da Cruz**: investigation. **R. Dyche Mullins**: Resources. **Nevan J. Krogan**: Conceptualization, supervision, and funding acquisition. **Robert W. Mahley**: Conceptualization, resources, writing - reviewing & editing, supervision, project administration, and funding acquisition. **Danielle L. Swaney**: Conceptualization, methodology, resources, writing - original draft, writing - reviewing & editing, visualization, supervision, project administration, and funding acquisition.

## Supplemental Figures

**Figure S1.**
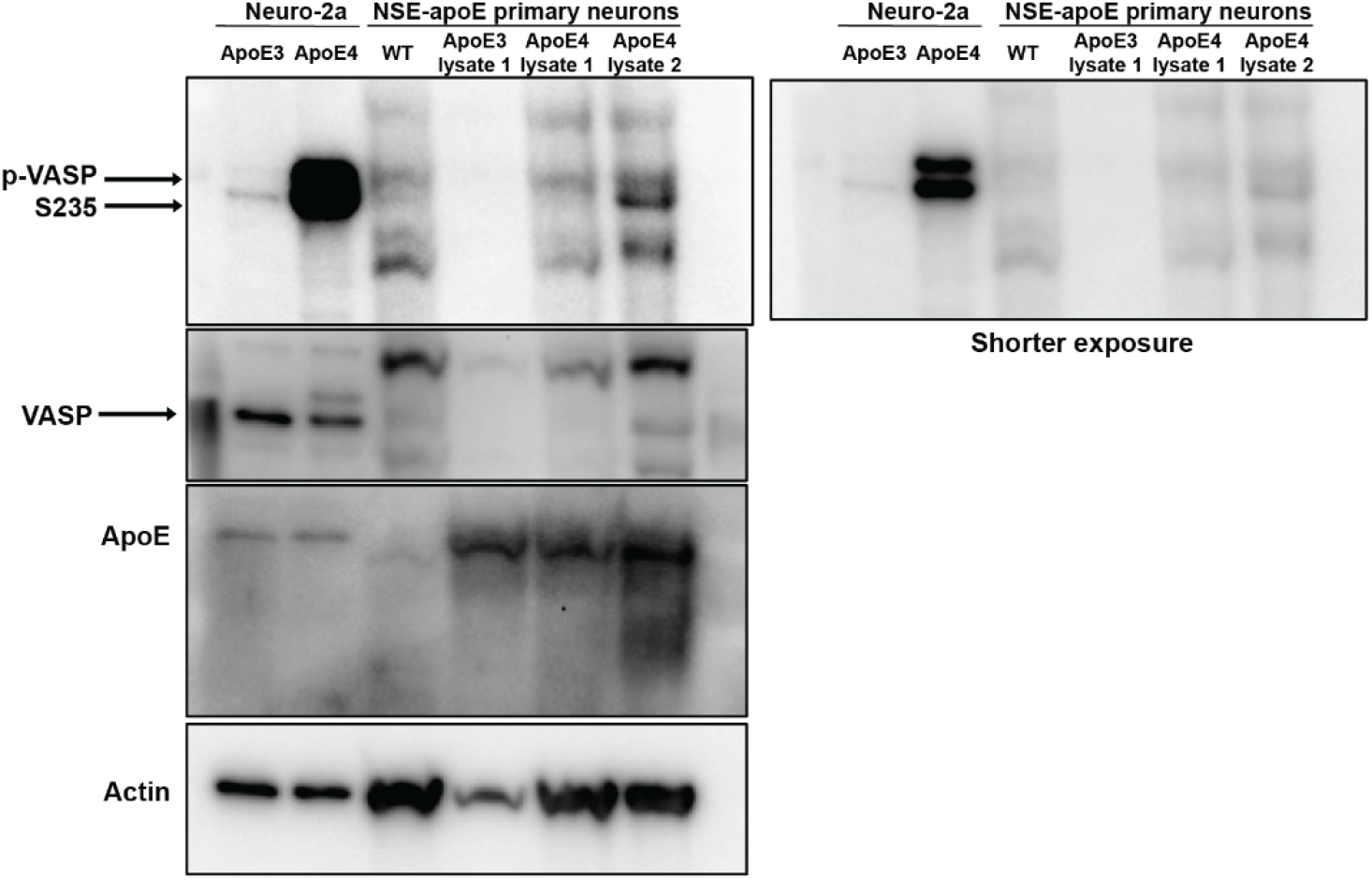
Immunoblotting of VASP 235 phosphorylation, VASP total protein, apoE, and actin in mouse primary neurons expressing apoE3 or apoE4. Actin serves as the loading control. Due to limited protein amounts lysates from different cultures were randomly pooled to provide sufficient protein amounts. This resulted in a single lysate for apoE3 neurons, and two lysates for apoE4 neurons. The right panel is a shorter exposure of the blot.

**Figure S2.**
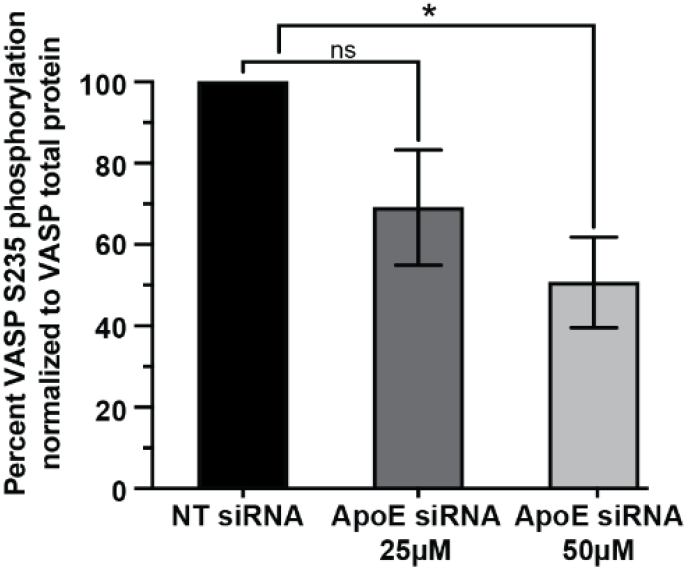
Quantification of VASP S235 phosphorylation levels upon apoE siRNA knockdown. Values are normalized to VASP protein levels, (n=2 biological replicates), ns:non-significant, *: p<0.05.

**Figure S3.**
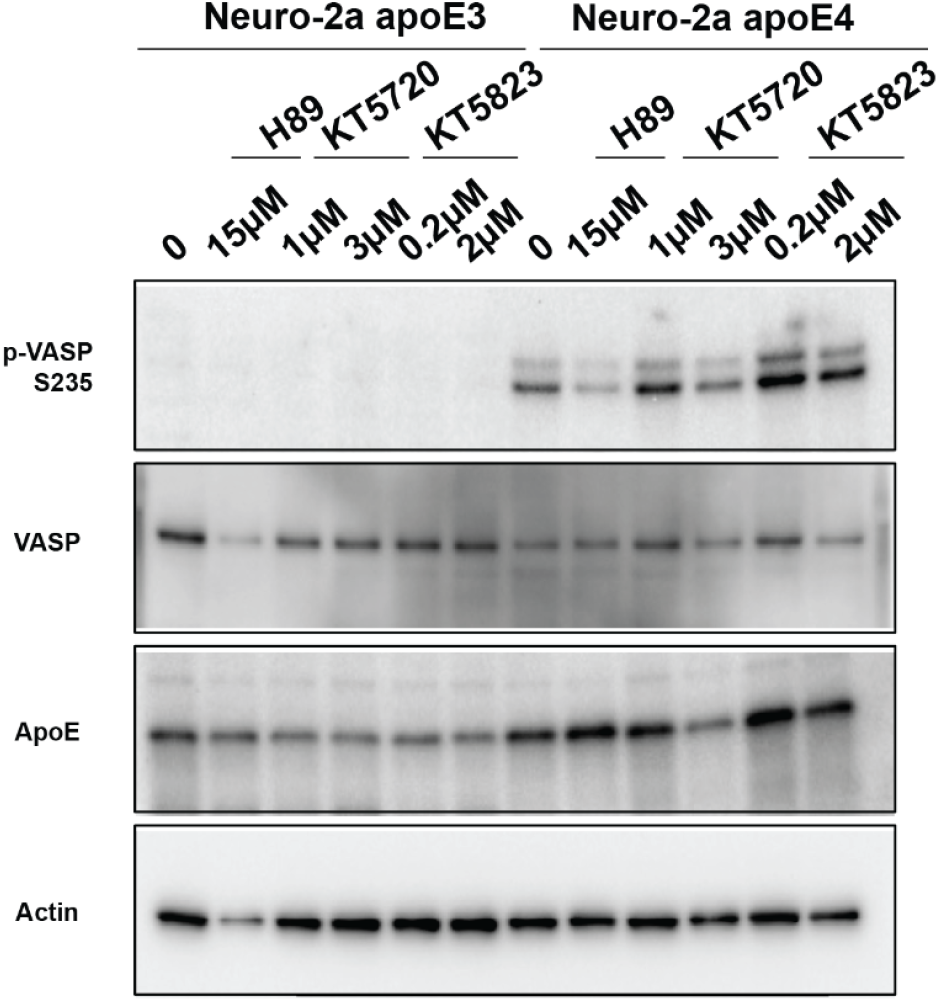
VASP S235 phosphorylation levels in apoE3 Neuro-2a and apoE4 Neuro-2a cells are determined by immunoblotting following treatment with PKA (KT5720) (1μM and 3μM) or PKG (KT5823) inhibitors. Cells were treated either with 1μM and 3μM of KT5720 or 0.2μM and 2μM of KT5823 for 24h.

**Figure S4.**
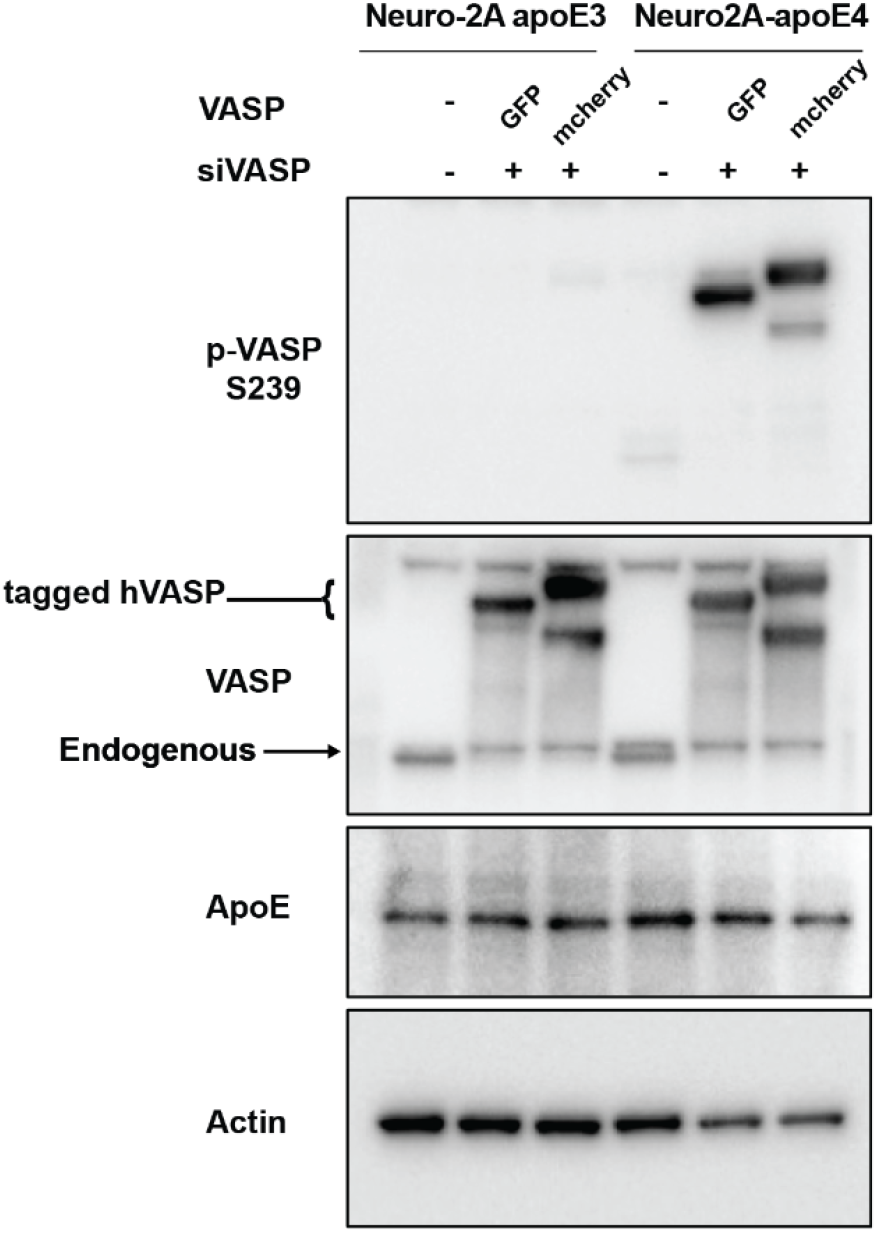
GFP and mcherry-tagged hVASP constructs were expressed in apoE3 and apoE4 Neuro-2a cells for 24h. Endogenous VASP was knocked down by VASPsiRNA. hVASP phosphorylation at S239 was validated by immunoblotting. Endogenous VASP is shown with arrows and hVASP is shown as tagged in brackets. Actin serves a loading control.

